# Surface topography is a context-dependent activator of TGF-β signaling in mesenchymal stem cells

**DOI:** 10.1101/2020.01.13.903195

**Authors:** Steven Vermeulen, Nadia Roumans, Floris Honig, Aurélie Carlier, Dennie G.A.J. Hebels, Aysegul Dede Eren, Peter ten Dijke, Aliaksei Vasilevich, Jan de Boer

## Abstract

We previously found that surface topographies induce the expression of the *Scxa* gene, encoding Scleraxis in tenocytes. Because *Scxa* is a TGF-β responsive gene, we investigated the link between mechanotransduction and TGF-β signaling. We discovered that mesenchymal stem cells exposed to both micro-topographies and TGF-β2 display synergistic induction of SMAD phosphorylation and transcription of the TGF-β target genes *SCXA*, *a-SMA*, and *SOX9*. Pharmacological perturbations revealed that Rho/ROCK/SRF signaling is required for this synergistic response. We further found an activation of the early response genes *SRF* and *EGR1* during the early adaptation phase on micro-topographies, which coincided with higher expression of the TGF-β type-II receptor gene. Of interest, PKC activators Prostratin and Ingenol-3, known for inducing actin reorganization and activation of serum response elements, were able to mimic the topography-induced TGF-β response. These findings provide novel insights into the convergence of mechanobiology and TGF-β signaling, which can lead to improved culture protocols and therapeutic applications.

## Introduction

Under physiological conditions, tissue homeostasis is maintained by the appropriate spatio-temporal responses of cells to environmental signals, such as secreted cytokines and transmembrane proteins from adjacent cells, but also by mechanical forces and changes in cell shape [1]. The latter is evident by the secretion and autocrine activity of insulin-like growth factor (IGF) by myocytes upon mechanical stimulation, a potent growth factor that induces muscle growth [2]. Similarly, tendon tissue homeostasis and growth depends on both mechanical forces [3] and transforming growth factor (TGF-β) signaling [4]. Also, the myofibroblastic state during tissue repair depends both on the mechanical characteristics of the matrix environment as well as the presence of TGF-β [5]. During embryonic development, mechanical forces are essential for proper morphogenesis in conjunction with biochemical signals, as shown by the spatial reorganization of TGF-β receptors upon cell confinement in gastruloids [6]. Mechanical forces can rapidly activate various intracellular signaling pathways [7]. Well-documented examples include the activation of the transcription factors yes-associated protein 1 (YAP), and tafazzin (TAZ) through stretching [8] and cell shape changes [9]. These transcription factors are essential for tissue homeostasis and embryogenic processes [10][11], such as osteogenesis [12], which is also influenced by bone morphogenetic protein 2 (BMP-2) [13]. Changes in actin dynamics influence serum response factor (SRF) activity through altered binding with co-transcription factors [14], leading to broad changes in physiological processes [15][16], including myofibroblast differentiation [17] which in turn can be regulated through TGF-β signaling [18]. How these mechanical and biochemical signals converge to drive cell behavior is poorly understood. Therefore, gaining novel insights in these mechanisms is vital for developing therapeutics in case of improper cell function and tissue engineering applications.

Since it is difficult to decouple the effects of biochemical and mechanical stimuli *in vivo*, essential insights are gained by *in vitro* experiments. Here, physical cues relayed through altered surface geometry can offer mechanical stimulation through changing cell shape. Cell geometry profoundly affects cell behavior, as shown by altering the lineage specification of stem cells [19][20][21]. Evidence exists that cell geometry also influences the biological effects of soluble factors. For example, adhesive islands alter the genomic response after tumor necrosis factor (TNF)α stimulation [22], and growth factor signaling from BMP-2 [23] or serum [24]. Also, cell confinement through micro-wells reduces the inflammatory response to lipopolysaccharide (LPS) in macrophages [25]. Of interest here is that surprisingly little is known how surface topography controls the cell’s response towards growth factor signaling. This is an important consideration since growth factors are involved in numerous biological processes, including differentiation [26] and maintenance of phenotypic identity [27]. Research involving cell stretching hints towards an interesting interplay between biomechanical forces and soluble factors. For example, cell stretching increases the sensitivity for soluble factors through altering receptor expression or activity [28][29][30]. In this manuscript, we provide new insights in this field by demonstrating that topographical cues alter the response of mesenchymal stem cells (MSCs) to TGF-β signaling.

## Results

### MSCs cultured on micro-topographies display altered actin dynamics and differential expression of cytoskeletal genes

An eye-catching characteristic of cells cultured on surface structures are profound changes in cell morphology [31][32]. We seeded adipose-derived MSCs on micro-topography PS-281 (Fig. 1A), and after 24 h observed changes in shape accompanied by a reduction of filamentous(F)-actin stress fibers compared to MSCs cultured on a flat surface (Fig. 1B-C). We previously assessed the transcriptome of bone marrow-derived MSCs on surface PS-281 relative to MSCs on flat control surfaces (manuscript in preparation) and produced a STRING protein-protein interaction network with 248 nodes and 1839 edges based on differentially expressed genes (DEGs) with a fold change > 1.5 and adjusted p-value < 0.05. The network includes early growth response gene 1 (*EGR1*) (Fig. 1D-E; 3.0 fold change), FBJ murine osteosarcoma viral oncogene homolog (*FOS*) (4.7 fold change), and *FOSB* (2.2 fold change), all of which are mechanosensitive transcription factors [33]. We next produced a volcano plot of genes with Gene Ontology (GO) term “*cytoskeleton organization*”, and noticed that many cytoskeleton related DEGs exhibit lower expression when cultured on surface PS-281, with a total of 48 DEGs downregulated and 11 DEGs upregulated (Fig. 1F and **Supplementary Table 1 and 2)**. Downregulated genes are involved in microtubule dynamics, e.g., stathmin 1 (*STMN1*; −1.6 fold change), an important cytoskeletal effector regulating microtubule dynamics [34], tubulin β class I (*TUBB;* −1.9 fold change), tubulin β 4B class IVb (*TUBB4B*; −1.8 fold change), tubulin α 1a (*TUBA1A*; −1.8 fold change), and tubulin β 6 class V (*TUBB6*; −1.6 fold change). Other genes such as actin γ-1 (*ACTG1*; −1.9 fold change) and tropomyosin 3 (*TPM3*; −1.7 fold change) form integral parts of the cytoskeleton [35]. Of interest, we also observed a slight downregulation of actin β (*ACTB*; −1.3 fold change). We further found a reduction in expression levels of genes associated with small GTPase rho signal transduction and subsequent cytoskeletal organization [36] **(Supplementary Fig. 1**). For example, we detected a downregulation of ezrin (*EZR*; −1.9 fold change), an actin-binding protein that acts as a linker between the actin cytoskeleton and plasma membrane proteins [37], and positively modulates rho signaling through the interaction with rho GDP dissociation inhibitors [38]. A downregulation was observed for diaphanous related formin 3 (*DIAPH3*; −1.5 fold change), which is required for F-actin stress fiber formation [39] and regulates SRF activity [40]. We also mention the downregulation of anillin actin-binding protein (*ANLN*; −1.7 fold change), which is important for cytoskeletal dynamics [41] and is involved in rho signaling [42]. The gene signature induced by surface PS-281 demonstrates that the cytoskeleton is under a lot of change 24h after cell seeding, which corresponds with the visual observed alterations in cytoskeleton architecture and cell geometry.

**Figure 1:**
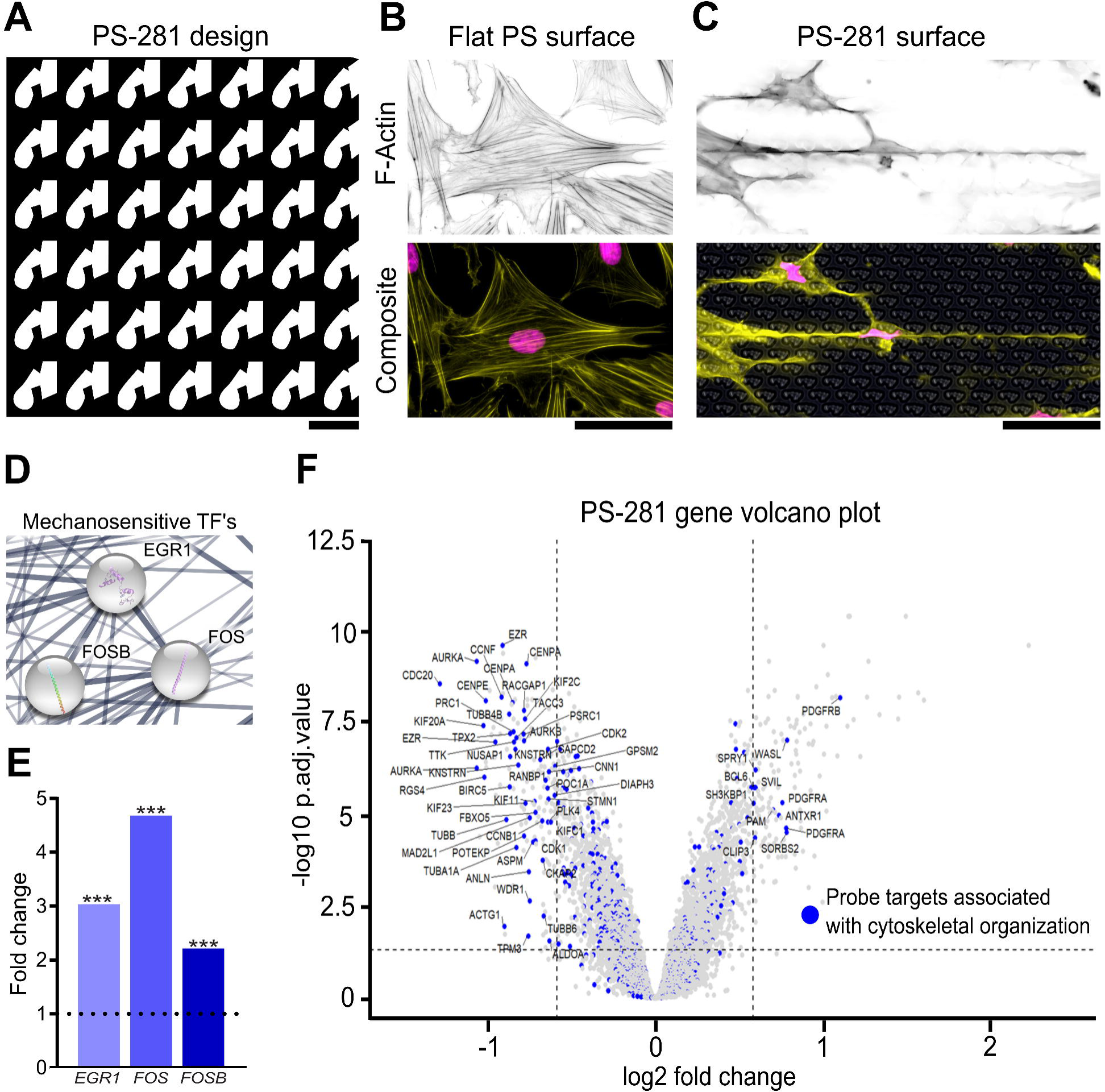
MSCs cultured on micro-topographies display altered actin organization and differential expression of cytoskeletal genes. **A)** *In silico* design of the PS-281 micro-topographical surface. Scale bar represent 10 μm. **B)** MSCs cultured on a regular PS flat surface exhibit spread morphological characteristics and a profound presence of F-actin stress fibers. **C)** MSCs cultured on surface PS-281 exhibit elongated and smaller nuclear and cellular characteristics, which coincides with a reduction of F-actin stress fibers. F-actin was immunolabeled with phalloidin (yellow) and the nucleus counterstained with Hoechst33342 (magenta). Scale bars represent 50 μm. **D)** Partial representation of a STRING gene network based on a microarray study of MSCs cultured on the PS-281 surface for 24h. The mechanosensitive transcription factors *EGR1*, *FOS*, and *FOSB* are represented here. **E)** *EGR1*, *FOS*, and *FOSB* expression levels have increased 2 to 5 fold on the PS-281 surface compared to a flat surface (*** P<0.001). **F)** Volcano plot representation of the PS-281 microarray data. Blue dots are probe targets associated with cytoskeletal organization. Majority of these DEGs exhibit lower expression levels compared to flat. DEG cut-off is determined at a 1.5 fold change and an adjusted P value of 0.05.

### Activation of early response genes is associated with early actin reorganization

Based on the observed increased expression of *FOS* and *EGR1* at 24h, we decided to investigate actin organization dynamics and the expression of early genes and proteins with a known relation to actin remodeling on surface PS-1018, which we previously discovered as a surface that induces Scleraxis (*Scx*) expression in tenocytes [43]. Furthermore, PS-1018 can be manufactured in a 100 mm dish format, thereby facilitating downstream experiments. Surface PS-1018 (Fig. 2A) induced cell elongation with a profound reduction in cell, and nuclear size, while reducing actin stress fibers (Fig. 2B), similar to surface PS-281. As early as 1h after cell seeding, we noted that cells on flat surfaces exhibited a diffuse actin pattern with a round cell morphology (Fig. 2C); however, MSCs adhering on the surface PS-1018 displayed different dynamics. Here, MSCs engulfed the micro-topographies with concentrated F-actin stress fiber formation on top of the structures (Fig. 2D), but a more diffuse actin pattern at the bottom and in between the structures (Fig. 2E). Quantification of F-actin levels showed significantly elevated F-actin levels at 1h and 2h on PS-1018, with levels peaking at 1h and dropping afterward **(Supplementary Fig. 2A-B**). To probe the early regulatory responses, we exposed MSCs cultured for 2h to flat and surface PS-1018 and analyzed the phospho-proteome by mass spectrometry **(Supplementary Fig. 3**). We detected increased levels of phosphorylated actin in cells cultured on surface PS-1018, indicating active cytoskeletal reorganization [44][45]. Also, we detected increased levels of phosphorylated adenylyl cyclase-associated protein 1 (CAP-1) and drebrin 1 (DBN1), which regulate actin dynamics [46][47]. These observations indicate that MSCs cultured on the micro-topography are subjected to dynamic cytoskeletal regulation, characterized by an early adaptation phase involving actin remodeling.

**Figure 2:**
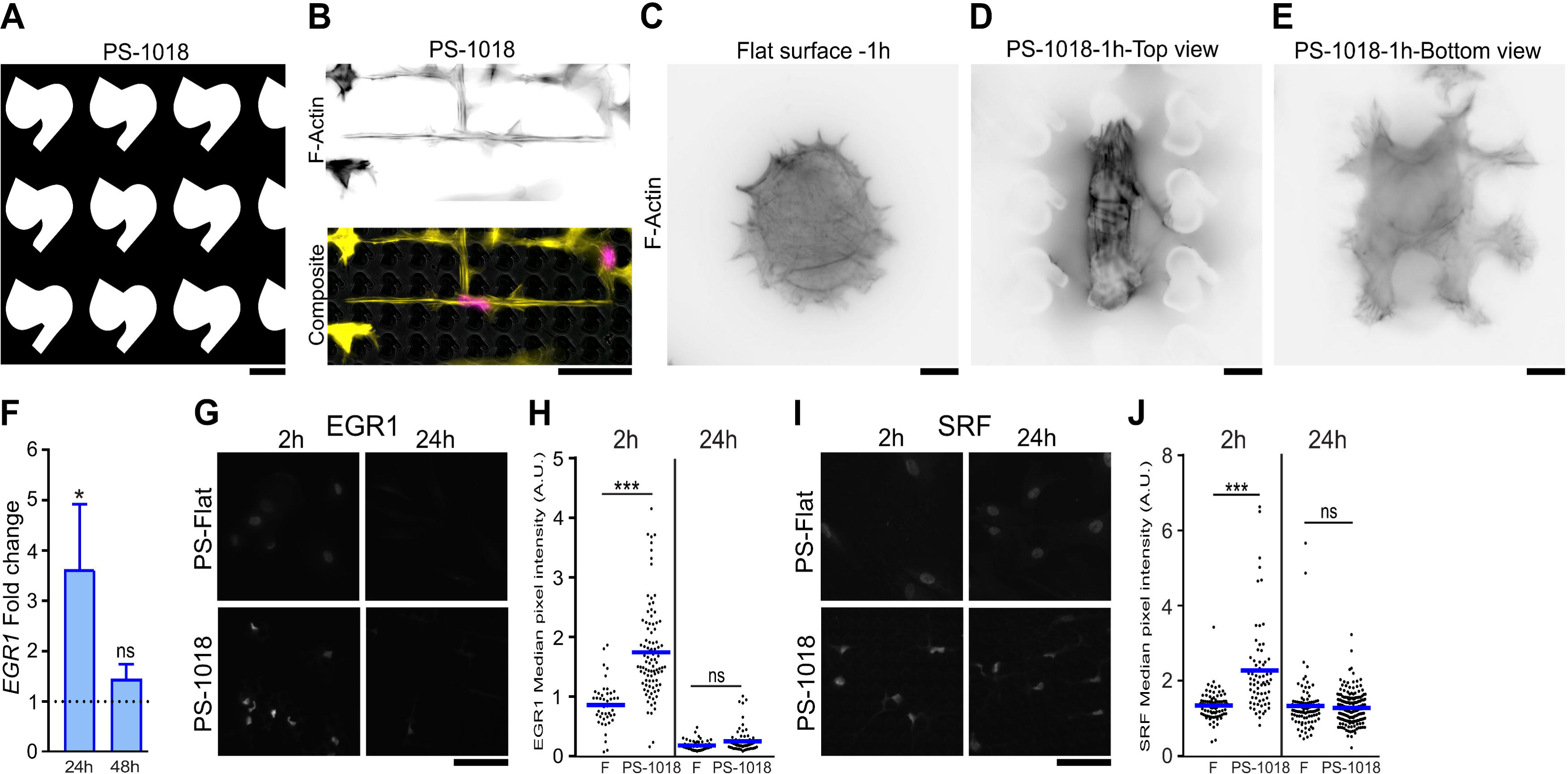
Micro-topographies elevate the early response genes SRF and EGR1 during the cells early adaptation phase. **A)** *In silico* design of the PS-1018 surface used in subsequent experiments. Scale bar represent 10 μm. **B)** Similar as the PS-281 surface, the PS-1018 surface elicits smaller, elongated morphological characteristics and a reduction in F-actin stress fibers. F-actin was immunolabeled by phalloidin (yellow) and the nucleus counterstained with Hoechst33342 (magenta). Scale bar represent 50 μm. **C)** Immunolabeling of F-actin after 1h of cell culture reveals that MSCs on a flat surface exhibit a round morphology with a diffuse F-actin pattern. **D-E)** Immunolabeling of F-actin of MSCs cultured for 1h on the PS-1018 surface reveals an increase in F-actin stress fibers concentrated on the upper part of the micro-topographical structures, while a diffuse pattern is observed at the bottom of the PS-1018 structures. Scale bar represent 10 μm. **F)** *EGR1* levels are elevated at 24h as measured through qPCR (* P<0.05), a similar observation as with the PS-281 microarray data. **G-H)** Immunolabeling of EGR1 demonstrates elevated intensities on the PS-1018 surface at 2h, with only a few cells on the PS-1018 surface showing elevated levels after 24h. Quantification of EGR1 fluorescent signal confirms the visual observation, with elevated levels measured at 2h (*** P<0.001), and no significant difference at 24h. Scale bar represent 100 μm. **I-J)** Immunolabeling of SRF demonstrates elevated levels on the PS-1018 surface at 2h (*** P<0.001), with no significant differences at 24h. Barplot represent the mean with error bars representing SEM. Scale bar represent 100 μm.

We next investigated genes and proteins, of which it is known that their expression or activity is influenced by actin. First, we observed a 3.6 fold increase in *EGR1* mRNA levels compared to flat at 24h and no significant difference at 48h (Fig. 2F). At earlier time points, we assessed EGR1 protein expression dynamics on both flat and PS-1018 (Fig. 2G). At 2h, we found increased EGR1 levels in the nucleus when MSCs are cultured on PS-1018 compared to flat (Fig 2H). Of interest, we measured a slight yet non-significant increase at 24h. This seemingly contradicts the qPCR and microarray observations on surface PS-281; however, we believe this caused by a subpopulation of cells with high EGR1 levels on the PS-1018 surface that skews the global *EGR1* levels measured on RNA level. Equally interesting was the observation that after 2h EGR1 levels on a flat surface were higher than on flat and the PS-1018 surface at 24h. We contribute this phenomenon to cell-seeding that induces a biomechanical response.

Next, we explored if we could detect alterations in SRF levels after 2h and 24h on PS-1018 (Fig. 2I). SRF is an important transcription factor that is associated with changes in actin dynamics [42] and known for inducing transcription of genes with serum response elements in its promoter, which includes FOS, EGR1, and SRF itself [48][49]. Similar to EGR1, high intensities of nuclear SRF were observed at 2h (Fig. 2J), followed by a slight yet non-significant decrease at 24h compared to the flat surface.

### Micro-topographical cues elevate TGF-βR-II and SCX levels in MSCs

EGR1 elevation on surface PS-1018 is interesting, considering that EGR1 is involved in the expression of the tendon-specific transcription factor scleraxis (*SCX*) [50][51], which in previous work was induced in tenocytes on micro-topographies [43]. It is unclear how EGR1 influences SCX but considering that SCX is upregulated in response to TGF-β [26], evidence exists that this is through increased expression of the TGF-β2 ligand [52] or the transforming growth factor-β type II receptor (TGF-βR-II) [53]. Browsing of the PS-281 transcriptomics data set for TGF-β signaling revealed five genes with a significant fold change of more than 1.5 associated with GO biological process “Response to TGF-β” (Fig. 3A). These include the previously mentioned *FOS* gene (4.6 fold change), known for participating with small mothers against decapentaplegic (SMAD) proteins to influence TGF-β signaling [54]. Also, an upregulation was observed for the TGF-β inducible genes Collagen-III (*COL-III*; 1.6 fold change) [55], prostate transmembrane protein androgen-induced 1 (*PMEPA1*; 1.5 fold change) [56], and matrix remodeling-associated protein 5 (*MXRA5*; 1.5 fold change) [57]. We also noted a downregulation of neuronal regeneration related protein (*NREP*; −1.5 fold change), which is related to an expression decrease of the growth factors TGF-β1 and TGF-β2 [58]. In addition, we noticed a increase in *SMAD7* expression (1.41 fold change), an antagonist of the TGF-β/SMAD pathway that functions as a negative feedback activator after TGF-β signaling [59]. We also mention a 2.81 fold change increase of *VCAM1*, which is TGF-β inducible [60], yet was not part of the GO list.

Of interest, we also noted (through multiple probes) an upregulation of *TGF-*β*R-II* (1.9 and 2.0 fold change), an essential component of the TGF-β/SMAD signaling pathway. These findings strengthen the hypothesis that micro-topographies sensitize MSCs for TGF-β related signaling. We confirmed increased *TGF-*β*R-II* expression on surface PS-1018 by qPCR after 8h (Fig. 3B; 1.3 fold change), and reaching a maximum at 24h (2.1 fold change). At 48h, no significant differences in *TGF*-β*R-II* levels were detected between PS-1018 and flat. It is known that mitogen-activated protein kinase kinase (MEK) inhibitors, such as U0216, can inhibit the activation of its downstream target *EGR1* [61][62], which we experimentally verified (**Supplementary Fig. 4**). Interestingly, U0216 also decreased *TGF-*β*R-II* expression on the PS-1018 surface.

**Figure 3:**
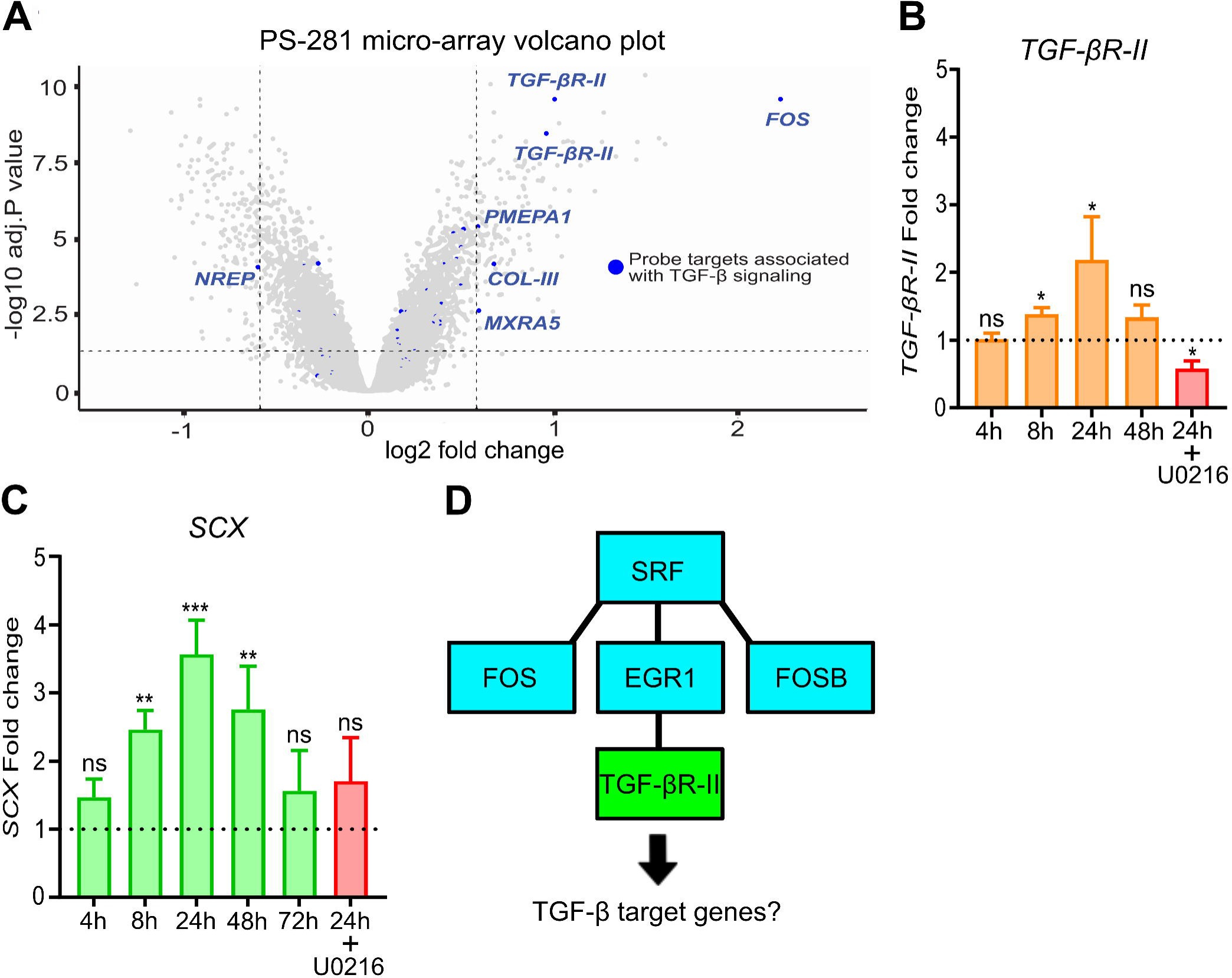
Micro-topographies induce elevated levels of *TGF-βR-II* and the TGF-β target gene *SCX*. **A)** Volcano plot of the PS-281 microarray with probe targets associated with TGF-β signaling represented in blue. DEG cut-off is determined at a 1.5 fold change and an adjusted P value of 0.05. Increased levels of *TGF-*β*R-II* are observed (1.9 and 2.0 fold increase; *** P<0.001). **B)** qPCR of MSCs cultured on the PS-1018 surface validates the PS-281 observation regarding *TGF-*β*R-II* expression, with elevated levels observed at 8h and 24h. The ERK inhibitor U0216 inhibits the topography induced *TGF-*β*R-II* expression (* P<0.05). **C)** The PS-1018 surface induces elevated levels of the TGF-β target gene *SCX.* Significant elevated levels were detected at 8h, 24h and 48h (** P<0.01), with a maximum expression at 24h. In addition, the inhibitor U0216 abolishes topography-induced *SCX* expression. **D)** Schematic representation of the mechanism involving the activation of SRF and EGR1 leading to an upregulation of *TGF-*β*R-II*. Barplots represent the mean with error bars representing SEM.

We further observed a 2.5 fold increase of the TGF-β inducible gene *SCX* after 8 hours, with maximum *SCX* levels after 24h (3.5 fold change), which decreased to 2.8 fold after 48 hours. After 72h, no significant *SCX* elevation was detected (Fig. 3C). We also found that U0216 reduced topography-induced *SCX* upregulation. These findings demonstrate that micro-topographies enhance *SCX* levels in MSCs, which could be guided by a general sensitization for TGF-β signaling through SRF, EGR1 (Fig. 3D), and TGF-βR-II.

### Surface topography and TGF-β2 synergistically induce TGF-β target genes

Given that *SCX* can be induced by both TGF-β2 [26] and topography [43], we investigated the combined effect of TGF-β2 and micro-topographies on TGF-β signaling. First, we measured SMAD2/3 phosphorylation (P-SMAD) as an immediate response to TGF-β receptor signaling and mediator of TGF-β target gene expression. 24h time after cell seeding, the timepoint with maximum *TGF-*β*R-II* expression, we exposed MSCs to TGF-β2 and fixed the cells 30 minutes after the treatment (Fig. 4A). Quantification of nuclear P-SMAD levels demonstrated that TGF-β2 treatment resulted in a 1.2 fold increase in nuclear P-SMAD levels compared to cells cultured on flat (Fig. 4B). Of interest, we observed that TGF-β2 treatment of MSCs grown on PS-1018 resulted in a 1.7-fold increase in nuclear P-SMAD levels compared to non-treated cells. Next, we isolated the RNA of MSCs cultured on flat or PS-1018, treated with and without TGF-β2 between 4h and seven days after cell seeding (Fig. 4C). The most striking and important observation we made is a synergistic induction of *SCX* expression at 24h. Whereas *SCX* levels on PS-1018 were 3.5 fold higher compared to a regular flat surface, TGF-β2 stimulation alone resulted in a 14.5 fold increase in *SCX* levels compared to flat. Of interest, is that the combined exposure of MSCs to PS-1018 and TGF-β2 induced *SCX* 39.9 fold. This synergy was already detected after 8 hours, although at lower levels, and was observed during the whole seven days culture period. It is interesting to note that *SCX* expression declines over time.

**Figure 4:**
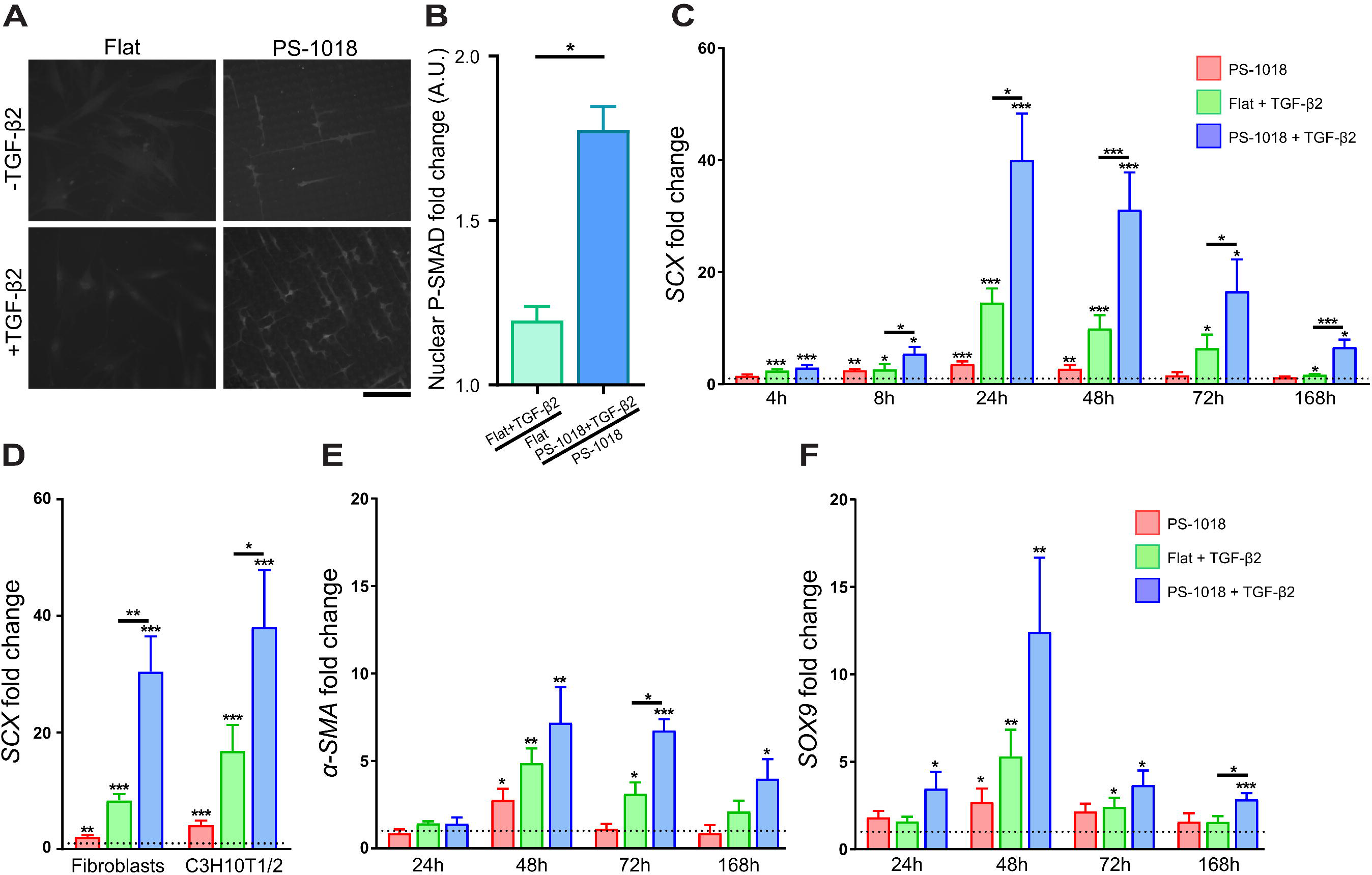
Micro-topographies positively modulate TGF-β signaling. **A-B)** P-SMAD immunolabeling revealed elevated nuclear distribution in MSCs cultured on the PS-1018 compared to flat when treated with TGF-β2 for 30 minutes. Scale bar represents 100 μm. **C)** TGF-β treatment induces elevated *SCX* expression already 4h after cell seeding, while PS-1018 induces an observable increase after 8h. Also at 8h, improved *SCX* levels are observed when combining micro-topographies with TGF-β2. Peak *SCX* levels are reached at 24h for all conditions. After 24h, overall *SCX* levels drop. Micro-topographies however retain their synergistically effect when combined with TGF-β2 at 48h, 72h and 7 days. **D)** Synergistic effect on SCX expression induced by the micro-topography can be reproduced with dermal fibroblasts and the mouse mesenchymal stem cell line C3H10T1/2. **E)** Improved α*-SMA* expression is observed when combining both PS-1018 and TGF-β2 at 72h and 7 days. **F)** Improved *SOX9* expression is observed when combing PS-1018 and TGF-β2 at 48h and 7 days (* P<0.05; ** P<0.01; *** P<0.001). Barplots represent the mean with error bars representing SEM.

The synergistic response to TGF-β2 on surface PS-1018 was not unique to MSCs. We induced a similar biological response in human dermal fibroblasts and C3H10T1/2 cells, a mouse mesenchymal-like cell line that is frequently utilized in differentiation studies (Fig. 4D). These findings demonstrate that the synergy between TGF-β signaling and surface topography is reproducible in multiple TGF-β2-response cell lines. Besides *SCX*, we also found a similar effect for other TGF-β responsive genes. Expression of α*-SMA*, a differentiation marker of smooth muscle cells and myofibroblasts [63], and the chondrogenic transcription factor *SOX9* [64] can be induced by TGF-β2, and display a synergistic effect when combined with micro-topographies (Fig 4E-F). These observations demonstrate that micro-topographies sensitize cells for TGF-β signaling.

### Rho/ROCK/SRF signaling is required for topography-induced TGF-β sensitization

We next set out to investigate the signaling events that occur between surface topography-induced actin-mediated signaling and transcriptional activation of TGF-β target genes, by investigating the synergistic effect in the presence of several small-molecule inhibitors of signal transduction. We confirmed the synergy in the presence of DMSO, the diluent of the inhibitors used in the rest of the study (Fig. 5A), and validated that *SCX* gene expression is indeed dependent on TGF-β receptor activation, using its inhibitor SB431542 [65] (Fig. 5B). Interestingly, this compound abolished *SCX* expression in MSCs on PS-1018 alone, which indicates that even without the addition of TGF-β2, TGF-β/SMAD signaling is occurring. This may hint at auto- or paracrine signaling elicited by the MSCs, or TGF-β originating from the serum media.

**Figure 5:**
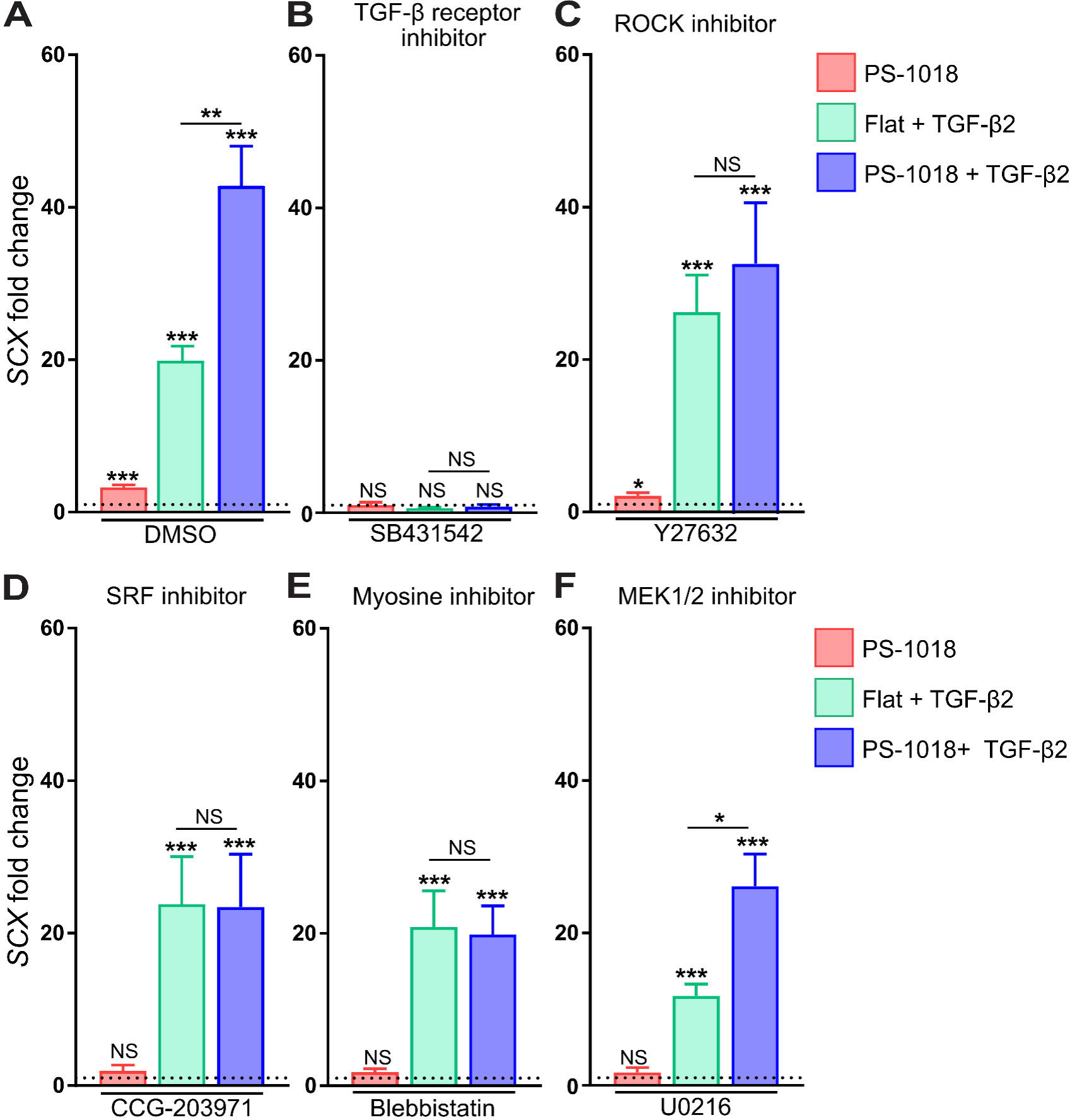
Pathway inhibitors reveal that micro-topographical induced *SCX* expression requires Rho/ROCK/SRF signaling. **A)** DMSO does not affect SCX levels in each condition. **B)** Inhibitors against the TGF-β receptor abolish *SCX* levels on all conditions including PS-1018, emphasizing the involvement of TGF-β signaling for micro-topographical induction of *SCX*. **C-E)** The Rho/ROCK/SRF inhibitors Y27632, CCG-209371, and blebbistatin abolishes the topography-induced effect on *SCX* expression. **F)** The MAPK/MEK inhibitor U0216 reduces overall SCX expression, yet does not reduce the synergistic effect on *SCX* expression (* P<0.05; *** P<0.001). Barplots represent the mean with error bars representing SEM.

Next, we studied the effect of the Rho-associated protein kinase (ROCK) inhibitor Y27632, because rho proteins influence numerous biological responses, including cell shape and actin cytoskeletal rearrangement [66] and are important for driving cell behavior of cells grown on physical cues. Y27632 did not affect the induction of *SCX* by TGF-β2 on flat control surfaces but did abolish the synergistic effect on the PS-1018 surface (Fig. 5C). Very similar results were obtained with CCG-203971, a compound which inhibits SRF/Myocardin Related Transcription Factor A (MRTF) gene transcription [67] (Fig. 5D) and blebbistatin (Fig. 5E), an inhibitor of non-muscle myosin-II [68] which prevents actin-myosin interaction resulting in subsequent disruption of actin dynamics [69]. Of interest, utilizing the MEK inhibitor U0216 did lower the level of SCX but did not abolish the synergistic effect (Fig. 5F), which indicates that the topography-induced mechanotransduction is not guided through MAPK signaling. These observations demonstrate that actin dynamics and the Rho/ROCK/SRF signaling pathway is necessary for micro-topographies to enhance the expression of *SCX*, but not for basal TGF-β2 activity.

### PKC activators mimic topography-induced mechanotransduction

Small molecules that inhibit actin-related signaling were able to block topographic induction of TGF-β2 signaling, so vice versa, it may be possible to mimic topographic mechanotransduction with small molecules that mimic actin-related signaling. To find these molecules, we searched the Connectivity Map, a compendium of more than one million gene expression profiles induced by small molecules and genetic perturbations, which is used for determining similarities in gene expression profiles between these perturbations [70]. We previously described that topography-induced TGF-β2 signaling coincides with high levels of F-actin and concomitant SRF signaling after 1h, but reduced F-actin and genes related to Rho/ROCK/SRF signaling after 24h. We reasoned that topography mimicking small molecules should, therefore, recapitulate the actin dynamics observed on topographies. The Connectivity Map only contains gene expression data from later time points, and we therefore retrieved small molecules using the gene expression fingerprint of cells in which β-actin, *SRF*, and *FOS* genes were knocked down as bait for the search (Fig. 6A-C), because this best reflects the lower Rho/ROCK/SRF signaling axis after 24 hours. Of interest, perturbagen class “Protein Kinase C (PKC) activators” was present in the lists of β-actin (*ACTB*), *SRF*, and *FOS* and thus resembles gene signatures associated with these knockdowns. Furthermore, in the list of PKC activators, “MEK inhibitors” were present with a negative score, which could indicate a positive involvement for EGR1 (Fig. 6D). Also for tubulin, of which we see a decrease of multiple isoforms in the microarray data, we see an association with PKC activators (**Supplementary Fig. 5**), suggesting that treatment of cells with PKC activators leads to similar gene expression profiles as when *ACTB*, *SRF*, and *FOS* are knocked-down.

**Figure 6:**
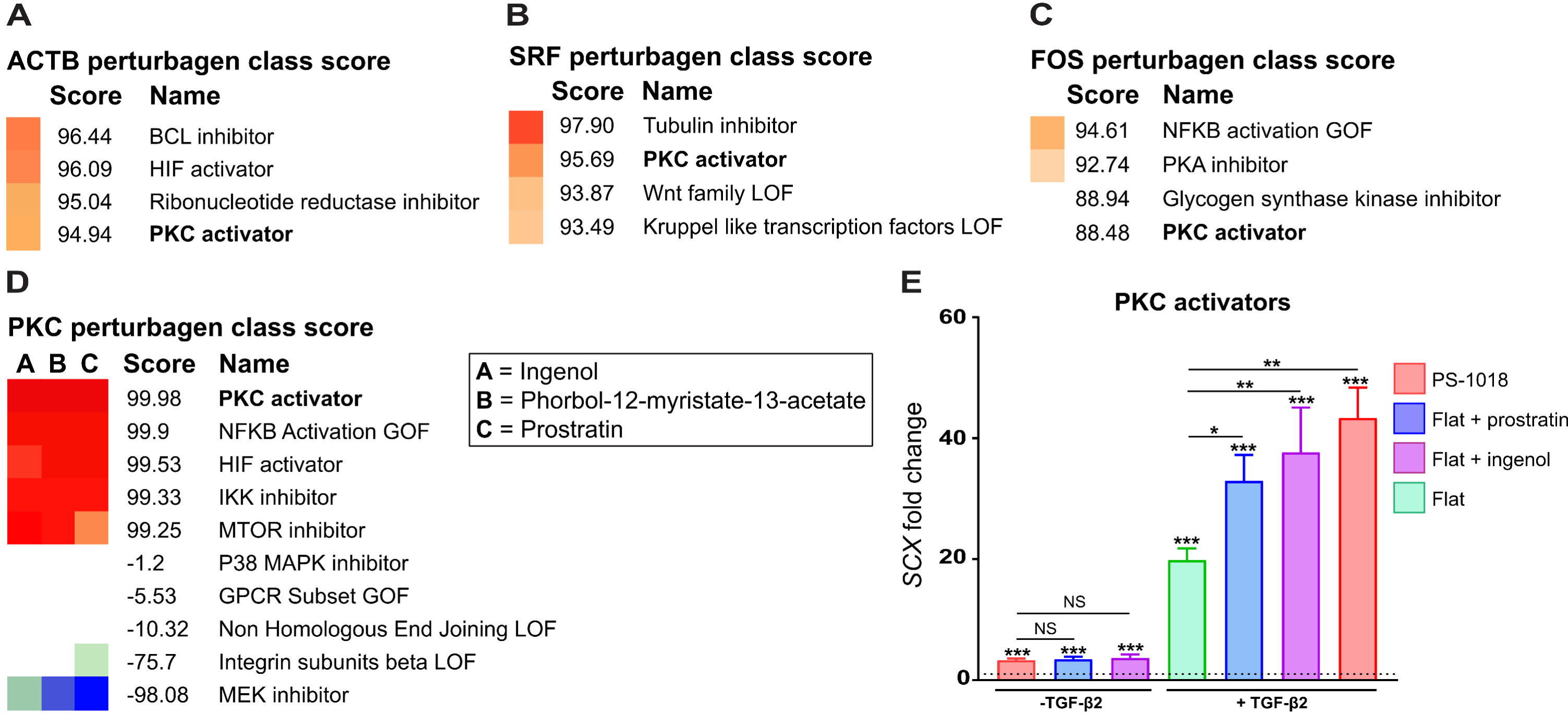
The connectivity map associates the gene signature of PKC activators with topography-induced mechanotransduction. **A)** *ACTB* knockdown gene signature corresponds with a PKC activator score of 94.94. **B)** *SRF* knockdown gene signature corresponds with a PKC activator score of 95.69. **C)** *FOS* knockdown gene signature corresponds with a PKC activator score of 88.48. **D)** PKC perturbagen class gene signature corresponds with a MEK inhibitor score of −98.08. **E)** Through qPCR, we found that the PKC activators prostratin and ingenol reproduce the micro-topographical effect on *SCX* expression, both with- and without the addition of TGF-β2 (* P<0.05; ** P<0.01; *** P<0.001).

PKC activators can induce actin reorganization [71], which eventually leads to a decrease in actin stress fibers [72]. Furthermore, they can activate *FOS* [73] and *EGR1* [74]. This makes this perturbagen class an interesting candidate for molecules that can mimic topography-induced TGF-β sensitization. We want to mention that in the β-actin knockdown gene fingerprint list, we found gene signature resemblances with other perturbagens such as cytochalasin-B, a microtubule inhibitor, and cytochalasin-D, an actin polymerization inhibitor (**Supplementary Fig. 6**). However, as shown with the actin polymerization inhibitor blebbistatin, this compound fails to recapitulate the synergistic effect with TGF-β2, since unlike PKC activators, no activation of early response genes is achieved. Also of interest is that an *EGR1* knockdown corresponds strongly with the gene signature of a *TGF-*β*R-II* knockdown, further emphasizing the relationship between these genes (**Supplementary Fig. 7**).

From the PKC activator component list (**Supplementary Fig. 8**), ingenol and prostratin were added to MSCs with and without TGF-β2, and *SCX* levels were measured after 24h. Without TGF-β2, ingenol and prostratin induced similar *SCX* levels as the PS-1018 surface (Fig. 6E). In the presence of TGF-β2, prostratin induced a 36.2 fold *SCX* expression compared to a regular flat surface without TGF-β2. Ingenol-3 induced a similar response, with slightly higher *SCX* levels (39.1 fold change) than prostratin. No significant differences between the topography and PKC activators were observed when adding TGF-β2. These findings demonstrate that PKC activators strikingly mimic the effect of the micro-topography in inducing *SCX* levels.

In previous work, we found that tenocytes rapidly lose their phenotypic characteristics in cell culture [43]. Cells transform from a spindle-shape towards a rounded morphology, which coincides with the formation of F-actin stress fibers. We were therefore interested in investigating if PKC activators could improve phenotypic characteristics in tenocyte culture. Therefore, we treated passage one tenocytes with- and without the PKC activators and TGF-β2 and found more profound spindle-shaped characteristics in confluent conditions when cells were treated with ingenol, both without and with TGF-β2 (Fig. 7A). To further investigate their morphological characteristics, we fixed cells 72h after adding the compounds and stained for F-actin, and SCX. Ingenol caused an apparent reduction in F-actin stress fibers and a decrease in F-actin intensity levels (Fig. 7B). Of interest is that TGF-β2 induces a more spread out morphology, which coincided with higher F-actin levels. This observation was also abolished by ingenol. We found that TGF-β2 was able to increase SCX levels, which could be further amplified by the ingenol treatment (Fig. 7C). These novel findings demonstrate that PKC activators replace the synergistic effects of the micro-topography both in the context of TGF-β induced MSC differentiation and phenotypic maintenance of tenocytes.

**Figure 7:**
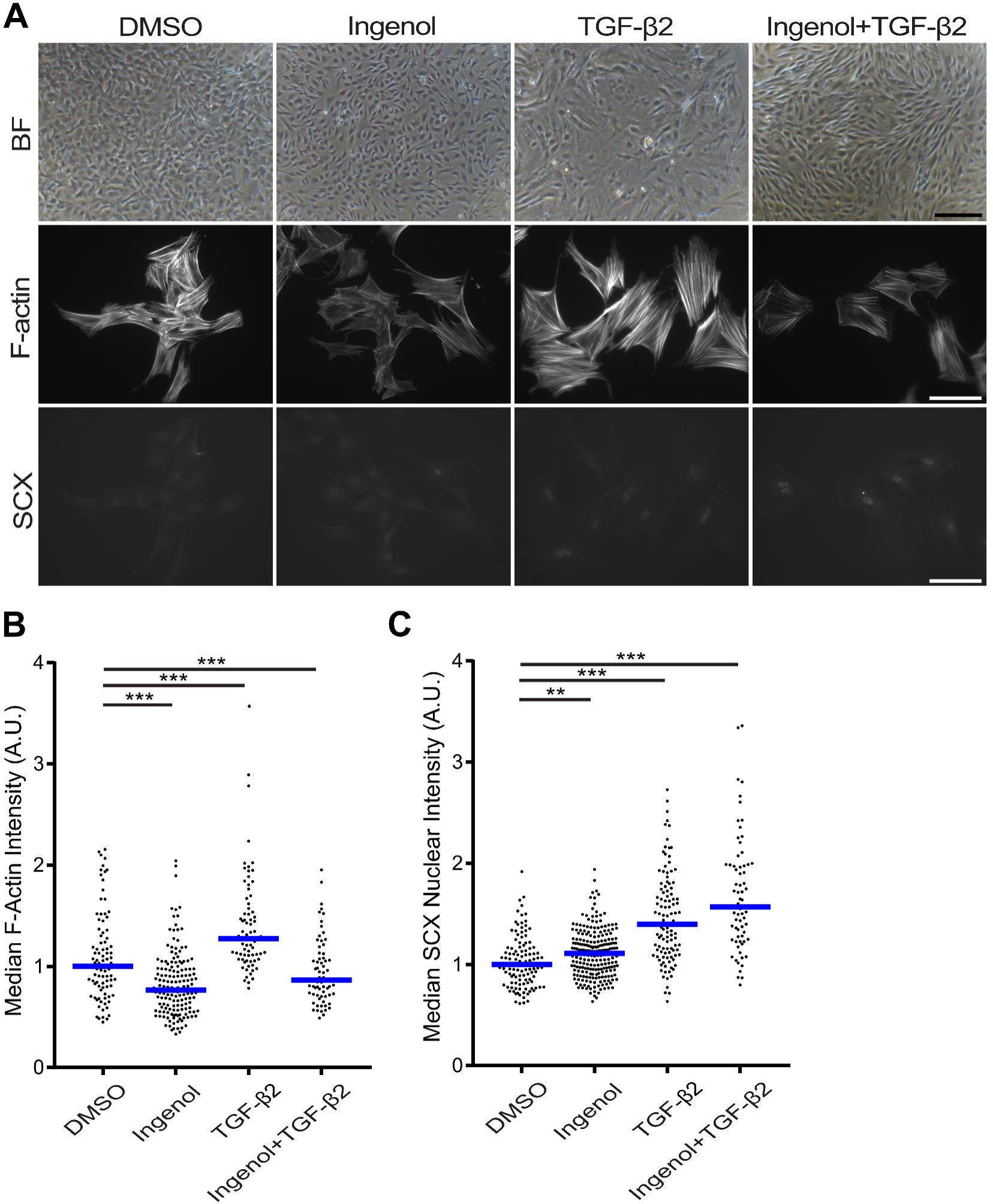
PKC activators reduce F-actin levels and elevate SCX protein levels in tenocytes. **A)** Ingenol-3 induces elongated cell characteristics in confluent conditions. Furthermore, reduced F-actin levels in tenocytes both with- and without a TGF-β2 treatment are observed when treated with ingenol. Similar was with MSCs, SCX levels increase by treating tenocytes with a PKC activator. Scale bar represent 100 μm. **B)** F-actin immunolabeling quantification reveals significant lower levels when tenocytes are treated with ingenol, both with- and without TGF-β2. **C)** SCX immunolabeling quantification reveals significant increased levels when cells are treated with ingenol and TGF-β2, with a most optimal effect when combining both perturbations (*** P<0.001).

## Discussion

We present evidence that TGF-β signaling is positively modulated through mechano-induced topographical cues. This work emphasizes the importance of recapitulating the crosstalk typically seen *in vivo* between mechanical and soluble cues in an *in vitro* culture setting. In addition, the results of this work put other research from the tissue-engineering field in a new perspective. BMP-2 is a growth factor that improves bone formation in a clinical setting, but only at un-physiologically high levels of the protein [75]. We hypothesize that BMP-2 is presented in the wrong mechanical context, and mechanical fine-tuning can lead towards greater efficacy. Research combining BMP-2 with mechanical loading indeed seems to indicate this [76]. Similarly, VEGF promotes angiogenesis, yet it also leads to vascular disruption [77]. Here it is plausible that the underlying mechanisms will be better controlled by implementing alternative stimulation as well. It is therefore interesting to reconsider growth factor-mediated cell differentiation and cell growth in the context of biomechanical conditions.

In an *in vitro* context, other studies involving geometrical-induced modulation of tenogenic gene expression do not consider the potential synergistic effects of growth factor signaling in their experimental setup [43][78][79][80][81][82]. An exception is a study documenting improved *SCX* expression in a dermal fibroblast culture on micro-grooves combined with TGF-β1 [83]. Besides micro-topographies, other physical cues such as scaffolds also enhance tenogenic characteristics of MSCs and progenitor cells [84][85], making this an interesting material design parameter for exploring if it might modulate TGF-β signaling. In this context, matrix stiffness can influence *SCX* expression [86], another physical environmental factor known for modulating TGF-β signaling [87][88][89]. These examples highlight the potential of future research involving biomaterials that investigate interactions with TGF-β signaling and other signaling cascades elicited by soluble cues.

In this study, we found mechanobiological signaling similarities with other experimental setups. For example, increased levels of EGR1 during the cell adaptation phase are also found early upon cell stretching together with increased levels of FOS [33]. EGR1 plays a crucial role in tenocyte mechanical signaling and can influence TGF-βR-II [53], SCX, and other tenogenic markers [51][50][52]. It is therefore not surprising that mechanical stimulation through stretching is studied extensively for inducing MSC differentiation [90][91], and tenogenic matrix deposition [92]. Furthermore, a clear link exists between mechanical forces and TGF-β/SMAD activation in numerous cell types [93][94][95]. SRF activity plays a central role in mediating actin dynamics [14][96], which is clearly altered and dynamically modulated by physical cues in this study. A plausible scenario is that SRF binds to serum response elements (SRE) in its promotor and those of FOS [48] and EGR1 [97], triggering the biological events we observe. This concept is further emphasized through the use of PKC activators, known to induce SRE activation [73], and thereby mimicking the effect of the topography. Although physical cues can regulate Rho/ROCK signaling [19], evidence points towards SRF as a subsequent and essential mediator in the observed biomechanical signaling events [22][24][25]. Also, experimental readouts that involve mechanical stretching can be blocked by inhibiting the Rho/ROCK/SRF pathway [98][99]. Related to this, the concept that SRF is involved in TGF-β signal transduction is not novel. Evidence exists that force-induced activation of α-SMA and subsequent fibrosis is blocked by inhibiting the Rho/ROCK pathway and subsequent SRF activity [17][100].

This research provides novel insights into how physical cues transmit mechanobiological signaling. Especially the dynamical nature of the signaling cascades elicited by the micro-topographies is fascinating. Parts of the mechanisms behind the phenomena we observe are however still unknown. For example, we do not rule out that FOS plays a role in TGF-β sensitization due to a binding affinity against SMAD2/3 [54]. Also, despite the apparent requirement of SRF for inducing TGF-β sensitivity, other transcription factors might play a role in activating SRE’s of EGR1 and FOS in conjunction with SRF [101]. More insights will need to be gained by investigating how much pathway overlap there is between physical cues, PKC activation, and mechanical stretching. The observation that PKC activators mimic topography-induced biomechanical stimulation is a fascinating concept since it extends the translational perspectives of our experiments. Small molecules inducing the same synergistic effect as micro-topographies could be introduced into a clinical context where mechanical loading events typically support tissue repair. For tendon tissue-engineering, this implies the potential to utilize PKC activators for stimulating a growth factor-mediated healing response, which might improve current clinical treatments [102].

## Conclusion

We demonstrate that micro-topographies influence TGF-β signaling in MSCs, leading to increased expression of the differentiation markers *SCX*, *SOX9*, and α*-SMA*. We connect the origin of this mechanobiological signaling with actin remodeling elicited by physical cues, which coincides with subsequent activation of early response genes associated with an upregulation of *TGF-*β*R-II*. Through extensive pathway analysis, we identify small molecule compounds that mimic the effect of the micro-topography. The results of this study can lead to improved protocols for the differentiation of MSCs or phenotypic maintenance of primary cells involving TGF-β signaling. Furthermore, the identification of small molecules that mimic mechanobiological stimulation might be applied in a clinical setting for replacing mechanical stimulation in conjunction with soluble cues.

## Materials and Methods

### Surface fabrication

A detailed description of the surface fabrication procedures is found elsewhere [50]. In short, the inverse pattern of the topographies was etched into a silicon wafer by directional reactive ion etching (DRIE). The same principle is applied for large surface areas containing one particular micro-topography, yet without walls to separate them. To facilitate demoulding procedures, the wafer was coated with a layer of Trichloro(1H,1H,2H,2H-perfluorooctyl)silane (FOTS, Sigma-Aldrich). Polydimethylsiloxane (PDMS; Down Corning) was cured on the silicon wafer to generate a positive mould and was subsequently used as a template to create a second negative mould in Ormostamp polymer (micro resist technology Gmbh), which serves as the mold for hot embossing the polystyrene (PS) films (Goodfellow). The hot embossing procedure was carried out at 140 °C for 5 min, and a pressure of 10 Bar, with a demoulding temperature of 90 °C. Before cell culture, the PS topographies were treated with oxygen plasma to improve cell adhesion for 30 s at 0.8 mbar, 50 sccm O2, and 100 W. Quality of the fabricated imprints was assessed using a Keyence VK-H1XM-131 profilometer.

### Cell Culture

Adipose-derived human mesenchymal stem cells (AD-hMSCs) and dermal fibroblasts (DF) used in this study were purchased from Lonza. AD-hMSCs and DF were isolated from a 42-year-old, and 27-year-old female respectively. C3H10T1/2 cells were purchased at ATCC. Basic medium for AD-hMSCs and DF consists of MEM Alpha GlutaMAX, no nucleosides (Gibco). For the culture of C3H10T1/2 cells, DMEM low glucose (Merck) was used as basic media. Basic media was supplemented with 10% v/v fetal bovine serum (FBS), 0.2 mM ascorbic-acid-2-phosphate (ASAP), and 10 U/mL Penicillin/Streptomycin. Cells were grown at 37 °C in a humid atmosphere at 5% CO_2_. For experimental purposes, cells at passage 3-4 were seeded at a density of 5000-10000 cells/cm^2^ on flat and the topographical surface. Human TGF-β2 (Peprotech), and mouse TGF-β2 (R&D Systems) were included in the media during cell seeding at a final concentration of 20 ng/ml. Pathway inhibitors blebbistatin, Y27632, CCG-203971, U2016, and SB431542 were purchased from Sigma-Aldrich and included in the media at a final concentration of 10 μM. For inhibitor studies, inhibitors were added to the cell media 1h prior and during cell seeding. PKC activators Ingenol, and Prostratin were purchased from Sigma-Aldrich, and added to the medium at a final concentration of 10 μM during cell seeding.

### Microarray study and pathways analysis

Bone marrow-derived human MSCs were seeded on topography PS-281 for 24 hours in basic medium at a density of 15,000 cells/cm2 in 24 well plates in three replicas. Total RNA was isolated using the Nucleospin RNA isolation kit (Macherey–Nagel). Then, from 100 ng of RNA, cRNA was synthesized using the Illumina TotalPrep RNA amplification kit. Both RNA and cRNA quality was verified on a Bioanalyzer 2100 (Agilent). Microarrays were performed using Illumina HT□12 v4 expression Beadchips. 750 ng of cRNA was hybridized on the array overnight, after which the array was washed and blocked. Then, through the addition of streptavidin Cy□3, a fluorescent signal was developed. Arrays were scanned on an Illumina Beadarray reader, after which raw intensity values were background corrected in BeadStudio (Illumina). Further data processing and statistical testing were performed using the online portal arrayanalysis.org. The probe□level raw intensity values were quantile normalized and transformed using variance stabilization (VSN). A detection threshold of 0.01 was used for reducing the number of false positives. A linear modelling approach with empirical Bayesian methods, as implemented in Limma package, was applied for differential expression analysis of the resulting probe□level expression values. P□values were corrected for multiple testing using the Benjamini and Hochberg method. Genes with a corrected p-value below 0.05 were considered differentially expressed.

To construct a gene network of the DEGs, we applied an online STRING analysis (https://string-db.org/). Only DEG with a fold change higher than 1.5 and an adjusted p-value lower than 0.05 were included in the list.

For identifying small molecules that mimic topography-induced pathways, we searched for relevant genes affected by the micro-topography in the connectivity map (https://clue.io/).

### Phospho proteomics study

#### In-liquid digestion

A total of 60 µg protein in 50 µl 50 mM ammonium bicarbonate (ABC) with 5 M urea was used. 5 µL of dithiothreitol (DTT) (20 mM final) was added and incubated at room temperature for 45 min. The proteins were alkylated by adding 6 µL of IAA solution (40 mM final). The reaction was taken place at room temperature for 45 min in the darkness. The alkylation was stopped by adding 10 µL of DTT solution (to consume any unreacted IAA) and incubated at room temperature for 45 min. For the protease digestion, 2 µg trypsin/lysC was added to the protein and incubated at 37ºC for 2 h. 200 µl of 50mM ABC was added to dilute the urea concentration and further incubate at 37ºC for 18 h. The digestion mix was centrifuge at 2x 10^3^g for 5 min and the supernatant collected.

Phospho-peptides were enriched by using TiO2 spin columns according the manufacturers protocol (Thermo Scientific). Samples were subsequently labelled with TMT isobaric mass tagging labelling reagent (10-plex; Thermo Scientific) according to the manufacturer’s protocol. In short, 60 μg of protein for each sample was used. The TMT labelling reagents were dissolved in 41 μl acetonitrile per vial. The reduced and alkylated samples and control were added to the TMT reagent vials. The reaction was incubated for 1 h at room temperature and quenched for 15 min by adding 8 μl of 5% hydroxylamine. Equal amounts of the samples and control were combined in a new vial and analysed by liquid chromatography-tandem mass spectrometry (LC-MS/MS).

#### Protein identification using LC-MS/MS

A nanoflow HPLC instrument (Dionex ultimate 3000) was coupled on-line to a Q Exactive (Thermo Scientific) with a nano-electrospray Flex ion source (Proxeon). The final concentration of the TMT labeled digest/peptide mixture was 0.33 μg/μl and 5 μl of this mixture was loaded onto a C18-reversed phase column (Thermo Scientific, Acclaim PepMap C18 column, 75-μm inner diameter x 15 cm, 2-μm particle size). The peptides were separated with a 90 min linear gradient of 4-68% buffer B (80% acetonitrile and 0.08% formic acid) at a flow rate of 300 nL/min.

MS data was acquired using a data-dependent top-10 method, dynamically choosing the most abundant precursor ions from the survey scan (280–1400 m/z) in positive mode. Survey scans were acquired at a resolution of 70 × 10^3^and a maximum injection time of 120 ms. Dynamic exclusion duration was 30 s. Isolation of precursors was performed with a 1.8 m/z window and a maximum injection time of 200 ms. Resolution for HCD spectra was set to 30,000 and the Normalized Collision Energy was 32 eV. The under-fill ratio was defined as 1.0%. The instrument was run with peptide recognition mode enabled, but exclusion of singly charged and charge states of more than five.

#### Database search

The MS data were searched using Proteome Discoverer 2.2 Sequest HT search engine (Thermo Scientific), against the UniProt human database. The false discovery rate (FDR) was set to 0.01 for proteins and peptides, which had to have a minimum length of 6 amino acids. The precursor mass tolerance was set at 10 parts per million (ppm) and the fragment tolerance at 0.02 Da. One miss-cleavage was tolerated, Phosphorylation of serienes, tyrosines and threonines as well as oxidation of methionine were set as a dynamic modification. Carbamidomethylation of cysteines, Tandem mass tag (TMT) reagent adducts (+229.162932 Da) on lysine and peptide amino termini were set as fixed modifications.

#### Immunocytochemistry

After cell culture, the cells were washed with phosphate buffered saline (PBS; Merck) and fixed with 4% (w⁄v) paraformaldehyde (Sigma-Aldrich) for 5 min at 37 °C. After a washing step, cells were permeabilized with 0.01% (v/v) Triton X-100 (Acros Organics) and blocked with goat serum (1:100; Sigma-Aldrich) in PBT (PBS + 0.02% Triton-X-100, 0.5% BSA) for 1h. Afterwards, cells were incubated with the primary antibody in PBT for 1h. After a washing step, cells were incubated with goat anti-rabbit secondary antibody conjugated to Alexa Fluor 647 (1:500; ThermoFisher), together with Phalloidin conjugated to Alexa Fluor 568 (1:500; ThermoFisher) in PBT for 1h. After washing, the nucleus was counterstained with Hoechst 33258 (1:1000; Sigma-Aldrich) for 10 min. After a subsequent washing step, surfaces were mounted on glass cover slides with mounting media (Dako). All washing steps were performed in triplicate with PBT. Primary antibodies used in this study are: anti-SCX antibody (1:200; Abcam; ab58655), anti-EGR1 antibody (1:200; ThermoFisher; T.126.1), anti-Phospho-Smad2/3 (1:200; Cell Signaling Technologies; 8828S) and anti-SRF antibody (1:200; Santa Cruz; sc-335).

#### Image analysis

Fixed samples were inverted and fluorescent images were acquired through the glass coverslip using a fully automated Nikon Eclipse Ti-U microscope in combination with an Andor Zyla 5.5 4MP camera. Fluorescent images were analyzed through CellProfiler 3.1.8 [103] applying custom-made pipelines. After illumination corrections, morphology of the nucleus was captured by the Otsu adaptive thresholding method applied on the Hoechst 33258 image channel. Subsequently, cell morphology was determined by applying propagation and Otsu adaptive thresholding on the Phalloidin image channel. Cells touching the border of the image were filtered out of the dataset. Missegmentation artifacts were removed by applying an arbitrary threshold on nuclei and cell size. After background correction, intensity values of the target of interest were calculated either inside the nuclear or cell area. The image software Fiji was used for image visualization [104]. Two-way ANOVA was used for determining statistical differences between conditions.

#### Quantitative polymerase chain reaction (qPCR)

After cell lysis by Trizol (Thermo Fisher), total RNA was isolated using the RNeasy Mini Kit (Qiagen) according to the manufacturer’s protocol. Reverse transcription was carried out using an iScript™ cDNA synthesis kit (Bio-Rad). Quantitative PCR was performed using the iQ™ SYBR® Green Supermix (Bio-Rad) in a CFX96 Touch Real-Time PCR Detection System (Bio-Rad). Relative expression was determined using the ∆∆Ct method. The geometric mean of the reference genes Glyceraldehyde 3-phosphate dehydrogenase (*GAPDH*) and TATA-Box Binding Protein (*TBP*) was applied for normalization. Primer sequences are listed in **Supplementary Table 3.** All data points represent an independent experiment, and qPCR data is based on at least three independent experiments. T-test statistics was applied to determine statistical differences between a condition and the flat control. Two-way ANOVA was used for determining statistical differences between conditions.

#### MSC osteogenic and adipogenic differentiation

To assess if the AD-MSCs were multipotent during the experiments, we investigated their potential for differentiation towards the osteogenic and adipogenic lineage. Differentiation of AD-hMSCs towards the osteogenic lineage was achieved by seeding AD-hMSCs at a density of 5 × 10^3^ cells/cm2. After 24h, medium was changed with either a control or mineralization medium. The mineralization media is basic media supplemented with 10% v/v fetal bovine serum (FBS), 10 U/mL Penicillin/Streptomycin, with 10 nM dexamethasone (Sigma) and 10 mM β-glycerol phosphate (Sigma), while control medium includes the same components except dexamethasone. Media was refreshed every 2-3 days and after 21 days, cells were fixed overnight at 4 °C with 4% formaldehyde (VWR) in PBS. Afterwards, osteogenesis was assessed through staining mineralized deposits with a 2% Alizarin Red solution (pH=4.2) for 2 min. Excess staining was washed off with demineralized water (**Supplementary Fig. 9A**).

Differentiation of AD-hMSCs towards the adipogenic lineage was achieved by seeding AD-hMSCs at a density of 15 × 10^3^ cells/cm2. After 24h, medium was replaced with either a control or adipogenic medium. The adipogenic media consist of basic media supplemented with 10% v/v fetal bovine serum (FBS), 10 U/mL Penicillin/Streptomycin, 0.5 mM 3-isobutyl-1-methylxanthine (Sigma), 0.2 mM indomethacin (Sigma), 10 μg/mL Insulin (Sigma), and 1 μM dexamethasone (Sigma). Control medium consisted solely of basic media with 10% v/v fetal bovine serum (FBS) and 10 U/mL Penicillin/Streptomycin. Media was refreshed every 2-3 days and after 21 days, cells were fixed overnight at 4 °C with 3.7% formaldehyde (VWR), 0.01 g/ml CaCl2.2H_2_O (Merck) in PBS. Afterwards, adipogenesis was assessed by rinsing the fixation solution with demineralized water, and subsequently incubating the substrates in a 60% (v/v) 2-propanol (VWR) for 5 min. Fat droplets were stained through a freshly filtered solution of 0.3 % (w/v) Oil Red O dissolved in 60% (v/v) 2-propanol (VWR). Afterwards, the substrates were washed in triplicate with demineralized water (**Supplementary Fig. 9B**).

## Acknowledgments

SV, NR, FH, DH, AD, AC, and JdB acknowledge the financial support of the Dutch province of Limburg. SV and AD are supported by the European Union’s Horizon 2020 Programme (H2020-MSCA-ITN-2015; Grant agreement 676338). PtD is supported by Cancer Genomics Center Netherlands (CGC.NL). AC gratefully acknowledges her VENI grant (number 15057) from the Dutch Science Foundation (NWO). We thank Freek Bouwman, Edwin Mariman, and Midory Thorikay for expert technical assistance.

---

**Supplementary Figure 1: STRING gene network representation illustrating RhoA interconnectedness with cytoskeletal genes.** Every dot represents a DEG in the gene network. Red dots are DEGs associated with the GO term “cytoskeletal organization”. For illustration purposes, RhoA was included in the network to demonstrate the strong relationship with cytoskeletal associated genes.

**Supplementary Figure 2: Early attachment of MSCs on micro-topographies is characterized by increased F-actin levels. A)** Immunolabeling by phalloidin allows visualizing F-actin in MSCs. During the early phase of cell attachment MSCs engulf the micro-topographies which coincides with increased F-actin levels. **B)** Quantifying F-actin of MSCs across multiple time points reveals that strongest intensities are detected on the PS-1018 surface 1h after cell seeding after which intensity levels declines (*** P<0.001). Scale bar represents 100 μm.

**Supplementary Figure 3: Phospho proteomics on MSCs after 1h of cell culture on PS-1018 reveals active cytoskeletal remodeling. A)** Phospho proteomics was able to detect both the phosphorylated and non-phosphorylated form of actin, adenylyl cyclase-associated protein 1, and isoform of drebrin 3. **B)** P-Proteomics reveals increased levels of phosphorylated actin and the actin associated proteins CAP1 and DBN1 (n=1).

**Supplementary Figure 4: MEK1/2 inhibitors downregulate EGR1 signaling. A-B)** Quantifying the EGR1 immunolabeling signal reveals that the MEK1/2 inhibitor reduces EGR1 levels 2h after cell seeding on PS-Flat (*** P<0.001).

**Supplementary Figure 5: Tubulin inhibitor perturbagen class score.** From this list, we notice the presence of PKC activators as inducing gene signatures similar as tubulin inhibitors.

**Supplementary Figure 6: ACTB compound class score.** PKC activators ingenol and phorbol-12-myristate-13-acetate are part of the list together with cytochalasin-d, an actin polymerization inhibitor, and cytochalisin-d, a microtubuline inhibitor. These inhibitors give rise to similar gene signature patterns in *ACTB* knockdown cell lines.

**Supplementary Figure 7: EGR1 genetic connectivity list.** From this list, it is clear that there is a strong connection between EGR1 and the TGF-βR-II.

**Supplementary Figure 8: PKC activator compound scores.** From this list, we selected ingenol and prostratin to assess their potential to replace micro-topographical induced signaling. MAPK and TGF-βR-II inhibitors induced opposite gene signatures.

**Supplementary Figure 9: Assessment of AD-MSC multipotency. A)** After osteogenic induction, mineralized calcium deposits were formed as visualized by Alizarin Red staining. **B)** After adipogenic induction, oil droplets were formed as visualized by Oil Red O staining.

