## Supplementary Table 1 for "Surface topography is a context-dependent activator of TGF-β signaling in mesenchymal stem cells"

| *Gene symbol* | *Entrez gene ID* | *Fold Change* | *adj.P.Val* | *Probe ID* |
| --- | --- | --- | --- | --- |
| SH3KBP1 | 30011 | 1.502874514 | 4.80E-06 | ILMN_1808501 |
| BCL6 | 604 | 1.505448135 | 1.79E-06 | ILMN_1737314 |
| CLIP3 | 25999 | 1.512117219 | 4.29E-05 | ILMN_1789733 |
| SVIL | 6840 | 1.512960691 | 1.79E-06 | ILMN_1671404 |
| SPRY1 | 10252 | 1.513235128 | 5.85E-07 | ILMN_2329914 |
| PAM | 5066 | 1.627279585 | 7.33E-06 | ILMN_2313901 |
| ANTXR1 | 84168 | 1.668567551 | 1.02E-05 | ILMN_1670379 |
| PDGFRA | 5156 | 1.689211807 | 4.66E-06 | ILMN_1681949 |
| PDGFRA | 5156 | 1.718928525 | 2.34E-05 | ILMN_2086470 |
| SORBS2 | 8470 | 1.722569828 | 2.97E-05 | ILMN_1716407 |
| WASL | 8976 | 1.723557946 | 9.22E-08 | ILMN_1666004 |
| PDGFRB | 5159 | 2.147994346 | 6.20E-09 | ILMN_1815057 |

**Supplementary Table 1**
