## Supplementary Table 2 for "Surface topography is a context-dependent activator of TGF-β signaling in mesenchymal stem cells"

| *Gene symbol* | *Entrez gene ID* | *Fold change* | *adj.P.Val* | *Probe ID* |
| --- | --- | --- | --- | --- |
| CDC20 | 991 | -2.435905142 | 2.64E-09 | ILMN_1663390 |
| AURKA | 6790 | -2.093044179 | 6.38E-10 | ILMN_1680955 |
| AURKA | 6790 | -2.090393096 | 5.19E-07 | ILMN_2357438 |
| KIF20A | 10112 | -2.033021327 | 3.66E-08 | ILMN_1695658 |
| RGS4 | 5999 | -2.0271293 | 9.28E-07 | ILMN_1758067 |
| CENPE | 1062 | -2.0154095 | 7.69E-09 | ILMN_1716279 |
| EZR | 7430 | -1.935043435 | 1.02E-07 | ILMN_3272378 |
| CCNF | 899 | -1.887069872 | 6.04E-09 | ILMN_1773119 |
| EZR | 7430 | -1.880320672 | 2.33E-10 | ILMN_1795937 |
| ACTG1 | 71 | -1.865228562 | 0.010975126 | ILMN_1704961 |
| TUBB | 203068 | -1.850651113 | 1.37E-05 | ILMN_1665583 |
| TUBB4B | 10383 | -1.827761605 | 1.73E-08 | ILMN_1780769 |
| BIRC5 | 332 | -1.825972531 | 1.73E-06 | ILMN_2349459 |
| NUSAP1 | 51203 | -1.822154177 | 2.54E-07 | ILMN_1726720 |
| TPX2 | 22974 | -1.8199839 | 6.05E-08 | ILMN_1796949 |
| CENPA | 1058 | -1.801457247 | 8.39E-09 | ILMN_1801257 |
| PRC1 | 9055 | -1.795554027 | 5.24E-08 | ILMN_1728934 |
| TTK | 7272 | -1.790995567 | 1.00E-07 | ILMN_1788166 |
| KNSTRN | 90417 | -1.780675801 | 1.58E-07 | ILMN_1652008 |
| TACC3 | 10460 | -1.773775699 | 7.87E-08 | ILMN_1724407 |
| TUBA1A | 7846 | -1.773233354 | 7.71E-05 | ILMN_1742981 |
| KNSTRN | 90417 | -1.761088335 | 4.32E-07 | ILMN_2141807 |
| PSRC1 | 84722 | -1.721294828 | 6.24E-08 | ILMN_1671843 |
| AURKB | 9212 | -1.719497282 | 9.54E-08 | ILMN_1684217 |
| POTEKP | 440915 | -1.719143514 | 3.75E-05 | ILMN_1814998 |
| RACGAP1 | 29127 | -1.718122473 | 1.38E-08 | ILMN_2077550 |
| KIF2C | 11004 | -1.714288742 | 2.36E-08 | ILMN_1685916 |
| KIF23 | 9493 | -1.710264427 | 4.73E-06 | ILMN_1811472 |
| CENPA | 1058 | -1.702362509 | 7.35E-10 | ILMN_2392472 |
| TPM3 | 7170 | -1.690793887 | 0.020426457 | ILMN_1697567 |
| ANLN | 54443 | -1.684261464 | 0.000354181 | ILMN_1739645 |
| MAD2L1 | 4085 | -1.68022029 | 1.18E-05 | ILMN_1777564 |
| WDR1 | 9948 | -1.679957565 | 0.002235003 | ILMN_1675844 |
| ASPM | 259266 | -1.654973658 | 5.48E-05 | ILMN_1815184 |
| KIF11 | 3832 | -1.6469069 | 4.29E-06 | ILMN_1794539 |
| FBXO5 | 26271 | -1.641126646 | 8.65E-06 | ILMN_1710676 |
| CDK1 | 983 | -1.639052483 | 5.07E-05 | ILMN_1747911 |
| SAPCD2 | 89958 | -1.606948727 | 3.06E-07 | ILMN_1702197 |
| CCNB1 | 891 | -1.594835947 | 1.47E-05 | ILMN_1712803 |
| CKAP2 | 26586 | -1.593481667 | 0.00017398 | ILMN_1674411 |
| TUBB6 | 84617 | -1.58657828 | 0.005790116 | ILMN_1702636 |
| RANBP1 | 5902 | -1.57294178 | 1.12E-06 | ILMN_1721457 |
| POC1A | 25886 | -1.562256691 | 1.82E-06 | ILMN_1780667 |
| CDK2 | 1017 | -1.558828044 | 1.58E-07 | ILMN_1665559 |
| PLK4 | 10733 | -1.558000535 | 1.54E-05 | ILMN_1789123 |
| STMN1 | 3925 | -1.553979956 | 3.62E-06 | ILMN_1657796 |
| CNN1 | 1264 | -1.551130861 | 6.53E-07 | ILMN_1810054 |
| ALDOA | 226 | -1.547705081 | 0.028255417 | ILMN_1736700 |
| KIFC1 | 3833 | -1.540762269 | 1.56E-05 | ILMN_2222008 |
| DIAPH3 | 81624 | -1.514809087 | 2.88E-06 | ILMN_1695226 |
| GPSM2 | 29899 | -1.51233378 | 4.75E-07 | ILMN_2139816 |
