## Supplementary Table 3 for "Surface topography is a context-dependent activator of TGF-β signaling in mesenchymal stem cells"

**Supplementary Table 5**

| ***Homo Sapiens*** | |
| --- | --- |
| **Primer Name** | **Sequence** |
| *SCX-F* | ACACCCAGCCCAAACAGA |
| *SCX-R* | GCGGTCCTTGCTCAACTTTC |
| *SOX9-F* | AAGACGCTGGGCAAGCTCTG |
| *SOX9-R* | GTAATCCGGGTGGTCCTTCTTG |
| *EGR-1-F* | AGCCCTACGAGCACCTGAC |
| *EGR-1-R* | GGGCAGTCGAGTGGTTTG |
| *α-SMA-F* | AAAAGACAGCTACGTGGGTGA |
| *α-SMA-R* | GCCATGTTCTATCGGGTACTTC |
| *TGFβR-II-F* | GTAGCTCTGATGAGTGCAATGAC |
| *TGFβR-II-R* | CAGATATGGCAACTCCCAGTG |
| *GAPDH-F* | TGTACCACCAACTGCTTAGC |
| *GAPDH-R* | GGCATGGACTGTGGTCATGAG |
| *TBP-F* | GAGCTGTGATGTGAAGTTTCC |
| *TBP-R* | TCTGGGTTTGATCATTCTGTAG |
| ***Mus musculus*** | |
| **Primer Name** | **Sequence** |
| *Scx-F* | AAACAGATCTGCACCTTCTG |
| *Scx-R* | TCAGATCAGGTCCAAAGTGG |
| *Gapdh-F* | CCTGGTCACCAGGGCTGC |
| *Gapdh-R* | CGCTCCTGGAAGATGGTGATG |
| *Tbp-F* | CTTCCTGCCACAATGTCACAG |
| *Tbp-R* | CCTTTCTCATGCTTGCTTCTCTG |
| **Rattus norvegicus** | |
| **Primer Name** | **Sequence** |
| *Scx-F* | GCACCTTCTGCCTCAGCAAC |
| *Scx-R* | TTCTGTCACGGTCTTTGCTCA |
| *Gapdh-F* | CCTGGTCACCAGGGCTGC |
| *Gapdh-R* | CGCTCCTGGAAGATGGTGATG |
| *Tbp-F* | TGGGATTGTACCACAGCTCCA |
| *Tbp-R* | CTCATGATGACTGCAGCAAACC |
